# Bone quality following peripubertal growth in a mouse model of transmasculine gender-affirming hormone therapy

**DOI:** 10.1101/2023.12.08.570840

**Authors:** Brandon W. Henry, Cynthia Dela Cruz, Robert W. Goulet, Bonnie T. Nolan, Conor Locke, Vasantha Padmanabhan, Molly B. Moravek, Ariella Shikanov, Megan L. Killian

## Abstract

During peri-puberty, bone growth and the attainment peak bone mass is driven predominantly by sex steroids. This is important when treating transgender and gender diverse youth, who have become increasingly present at pediatric clinics. Analogues of gonadotropin-releasing hormone (GnRH) are commonly prescribed to transgender and gender diverse youth prior to starting gender-affirming hormone therapy (GAHT). However, the impact of GnRH agonists on long bones with the addition of GAHT is relatively unknown. To explore this, we developed a trans-masculine model by introducing either GnRHa or vehicle treatment to female-born mice at a pre-pubertal age. This treatment was followed by male GAHT (testosterone, T) or control treatment three weeks later. Six weeks after T therapy, bone quality was compared between four treatment groups: Control (vehicle only), GnRHa-only, GnRHa + T, and T-only. Bone length/size, bone shape, mechanical properties, and trabecular morphology were modulated by GAHT. Independent of GnRHa administration, mice treated with T had shorter femurs, larger trabecular volume and increased trabecular number, higher trabecular bone mineral density, and wider superstructures on the surface of bone (e.g., third trochanters) when compared to control or GnRHa-only mice. In conclusion, prolonged treatment of GnRHa with subsequent GAHT treatment directly affect the composition, parameters, and morphology of the developing long bone. These findings provide insight to help guide clinical approaches to care for transgender and gender diverse youth.

## Introduction

Sex steroids play a crucial role in the attainment and maintenance of peak bone density (1). While sex steroids affect individuals during all ages of life, their effects are at their highest and most active during peri-pubertal growth affecting body composition and peak bone mass.(2– 4). In peri-pubescent boys, testosterone (T) stimulates periosteal apposition, leading to increased bone width compared with peri-pubescent girls, despite comparable cortical thickness between sexes (2). Estrogen (E) plays a major regulatory role in bone metabolism in both men and women, especially post-puberty, as it maintains bone remodeling and homeostasis (2). During development, estrogen promotes closure of the epiphysial plate during puberty; testosterone can induce similar effects via conversion to estrogen by aromatase (5). The release of sex steroids is regulated by signaling within the hypothalamo-pituitary gonadal (HPG)-axis. Gonadotropin-releasing hormone (GnRH) is secreted by the hypothalamus and stimulates the release of pituitary gonadotoprins that activate gonadal production of sex steroids like T and E. Inhibition of the HPG-axis leads to a hypoestrogenic state, limiting the role of estrogen in bone homeostasis (6,7).

Puberty suppression in transgender and gender diverse adolescents has been used historically to lessen the psychological burden of gender dysmorphia by making physical secondary sex characteristics more congruent with the experienced gender (8–10). One way this is accomplished is through administration of a GnRH agonist (GnRHa). Within the HPG-axis GnRHa will suppress the production and section of luteinizing hormone (LH) and follicle stimulating hormone (FSH), which results in arrested ovarian folliculogenesis and steroidogenesis. Adolescent patients at Tanner Stage 2-3, with persistent feelings of gender dysphoria that worsens at the onset of puberty, are eligible for GnRHa treatment according to International Guidelines (8,11,12). GnRHa can be prescribed as early as 10-11 yrs for trans boys and 11-12yrs for trans girls. Around 15-16 yrs of age, T or E can be used as gender affirming hormone therapy (GAHT) for full hormonal transitioning in consenting sufficiently capacitated patients (8,11,12). While much of what we know about GAHT in humans has focused on cardiovascular health and metabolic risk (13), the effect of delayed or prolonged pubertal suppression transitioned into GAHT on bone health is underappreciated. Previously, a decrease in areal bone mineral density (aBMD) Z-scores were reported after GnRHa administration, which were partially restored after GAHT (14).

While the effect of sex hormones on bone morphology and properties have been heavily studied, the impact of induction of the hypoestrogenic state in combination with T treatment in peripubertal mice such as that used in GAHT has not been explored extensively. In a model like ours, Dubois et al reports that GnRHa administration reduced bone mass acquisition, and bone strength in female to male (transmasculine) mice (8). This effect was also reversed by administration of T, achieving parameters equal to control female mice and in some cases male control (8). In this study, we examined the effect of GnRHA treatment with and without testosterone treatment on long bones in mice during peripubertal skeletal growth. Specifically, we measured femoral bone quality in adult female-born mice subjected to pubertal suppression followed by testosterone treatment to assess changes in the morphology of trochanteric, cortical, trabecular bone and its mechanical properties.

## Methods and Materials

### Experimental Design

C57BL/6N female mice (N=28) and age-matched male mice (N= 3) were used in this study. Female mice were assigned to one of four experimental groups to mimic gender-affirming hormone therapy (GAHT) in human adolescents, and treatment was initiated at postnatal age 26 days (P26), which is approximately the time female mice enter puberty (Figure 1) (15).

**Figure 1.**
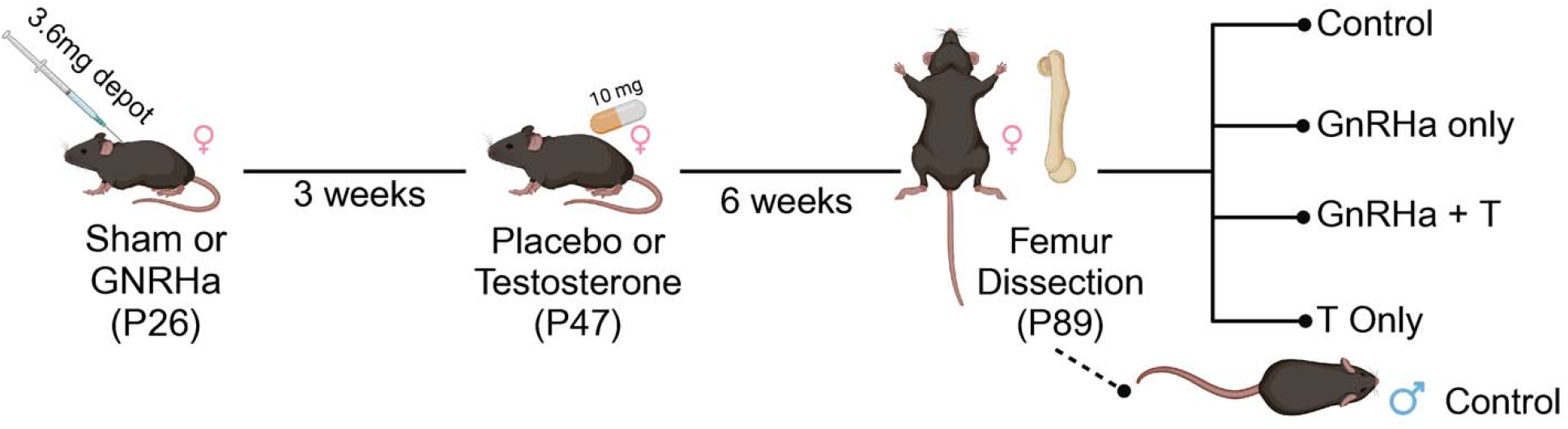
Summary of experimental design. At P26, C57BL/6N female mice were treated with GnRHa or sham. After 3 weeks, mice were supplemented with Testosterone (n=14) or placebo (n=14). Animals were euthanized at P89, then compared to control group (Sham + Placebo) and untreated P89 male mice.

Experimental groups included controls (sham surgery with empty silastic tubing implant; n=7), GnRHa-only treatment (for pubertal suppression; n=7), T-only treatment (for male transition; n=7), and GnRHa + T treatment (combined; n=7). For the GnRHa and GnRHa + T groups, mice were subcutaneously implanted with GnRHa (3.6mg) depot at P26. Control and T-only groups underwent a sham surgery with no implantation. At 3 weeks post-implantation or sham surgery, mice were subcutaneously implanted with silastic tubing containing testosterone (10 mg) or empty silastic tubing (control and GnRHa-only group). Six weeks after T-implantation, all animals were euthanized (Figure 1). Body weights were recorded at time of euthanasia and one femur from each mouse was dissected and fixed in 4% cold paraformaldehyde for 24-48hr then stored in 70% ethanol.

### Nano-computed tomography

Fixed femurs were imaged using nano-computed tomography (nanoCT; nanotom-m, phoenix|x-ray, Wunstorf, Germany) to obtain high resolution three-dimensional images of the femurs (parameters: 7.9 mm voxel size; 80kV; 400mA; 0.762mm aluminum filter; average parameter of 3; and integration time of 500ms). Calibration phantoms containing air, water, and a hydroxyapatite mimic (1.69mg/cc; Gammex, Middleton, WI, USA) were included in each scan. Full image volumes were reconstructed using datos|x software (GE, Inspection Technologies, LP, Skaneateles, NY, USA), and gray values were converted to Hounsfield units using the calibration phantoms. Dragonfly software (version 2021.1; Object Research Systems Inc, Montreal, Canada) was used to create a volume of interest (VOI) and a region of interest (ROI) of each imaged femur (Figure 2). A validated fully convolutional neural network (FCNN) was used to segment bone from background (DICE coefficient = 0.983 ± 0.016). A second validated FCNN was used to segment cortical from trabecular bone (Dice coefficient = 0.977 ± 0.023) to generate cortical, trabecular, third trochanter sub-volumes for structural quantification (16). The segmented image files were visually inspected by an expert (B.H.) for thresholding quality and cortical-trabecular segmentation (Figure 2). Erroneously identified voxels were manually corrected (16).

**Figure 2.**
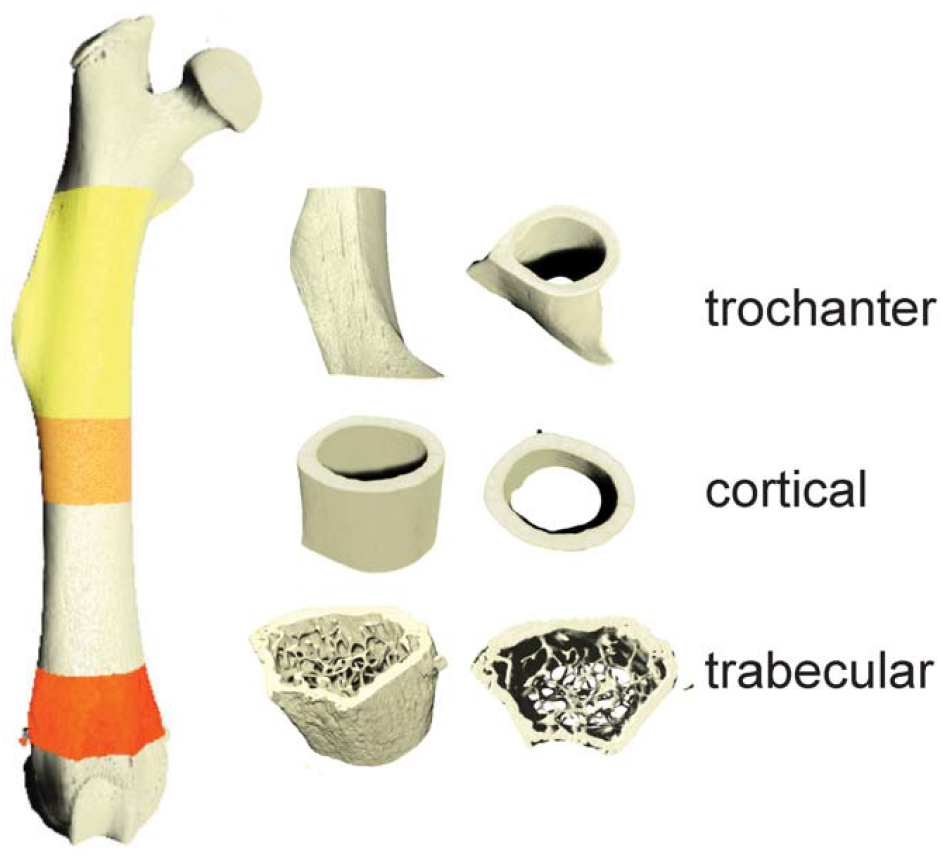
High resolution nanoCT was used to generate volumetric images of femurs (7.9mm voxel size). Analyses were performed on regions of interests including third trochanter, cortical, and trabecular regions.

All images were rotated to align the femoral long axis with the image Z-axis and the third trochanter with the X-axis for analysis. The boundary between cortical (Ct.) and trabecular (Tb.) bone was defined by prior studies (17,18). A boundary formula of the third trochanter was modeled after the cortical and trabecular bone boundary. We defined the start of the third trochanter ROI as the proximal boundary of the cortical bone where the periosteal surface starts to extend outward. The height of the third trochanter (in the long-axis Z dimension) was created by taking the 15% percent of total bone length. Third trochanter diameter was defined as the lateral margin of the third trochanter to the medial border to the femoral shaft (Figure 3A’) (19). Standard cortical and trabecular morphometric properties were calculated in each VOI (BV = bone volume; TV = total volume; I_max_ = maximum moment of inertia; Th = thickness, Ar = area, Ma = marrow, Pm = perimeter, BMD = bone mineral density; BMC = bone mineral content; Sp.= spacing, N = number). Data were compared using a two-way ANOVA correcting for multiple comparisons (Prism v9.0; Graphpad, LaJolla, CA). Data were presented as mean ± standard deviation.

**Figure 3.**
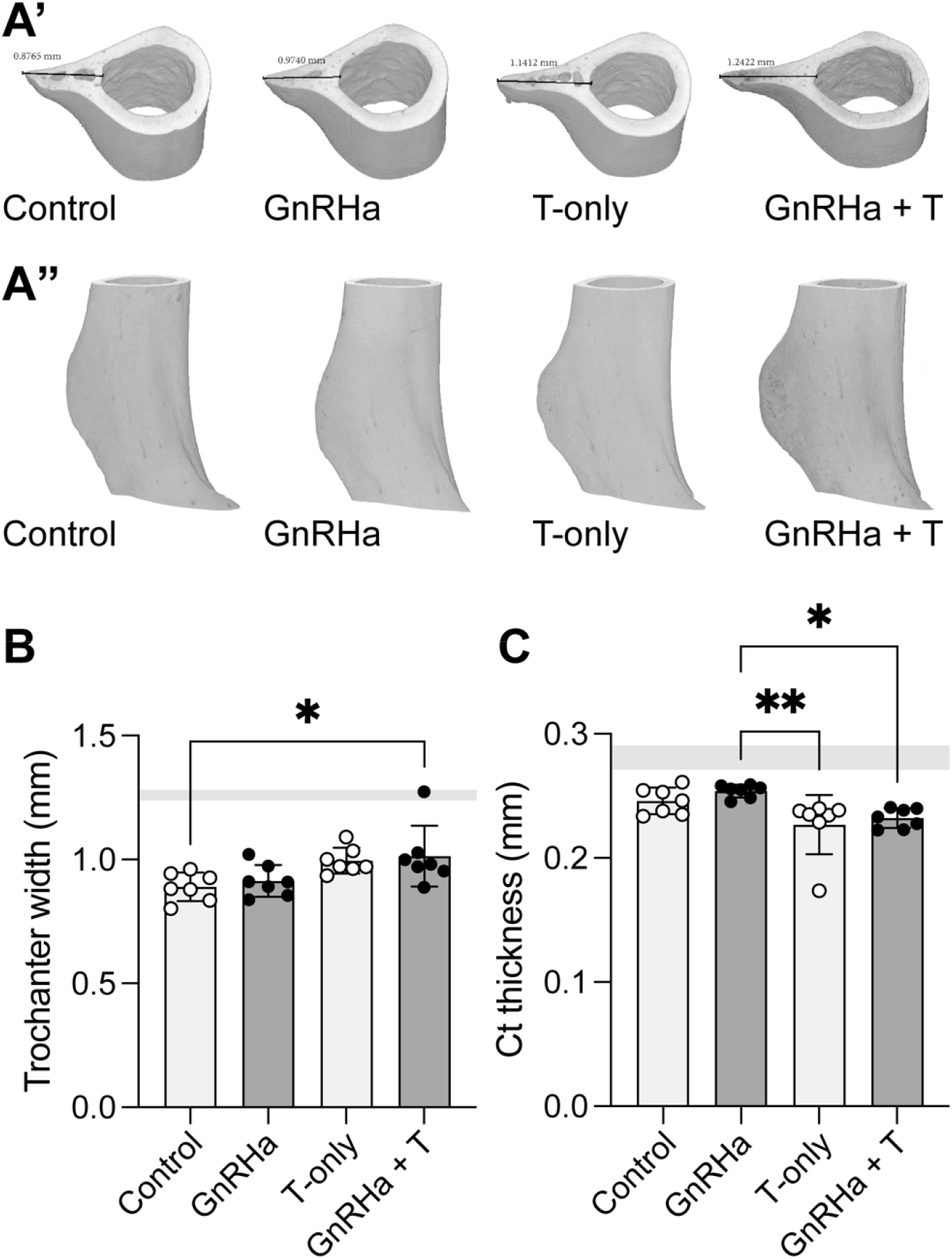
Third trochanter bone parameters in mice prepubertally treated with gonadotropin-releasing hormone analogues (GnRHa) and supplemented with gender-affirming hormones (GAH). (A’) Transverse 3D reconstruction of femoral third trochanteric bone from P89 female mice displaying peak third trochanteric length measured in mm. (A”) Full 3D reconstruction of femoral third trochanteric bone region of interest from adult female mice. (B) Third trochanter width was larger in groups treated with T compared to control. (C) Cortical bone was thinner in female mice treated with T compared to control or GnRHa only. Data are presented as biological replicates (individual dots) and mean +/-standard deviation. Gray horizontal bars represent range for age-matched male control mice, n=3.

### Mechanical Testing

Contralateral femurs (n = 6 females per treatment; n=3 males) were loaded to failure in 4-point bending with a span length of 6.5mm and 2.2mm for the top and bottom fixtures, respectively (MTS Systems, 858 Mini Bionix II) (20,21). Fixtures were positioned at mid-femur with the posterior side of the femur under tension and samples were kept hydrated using PBS. Femur length was measured using calipers from the greater trochanter to the lateral femoral condyle. Femurs were loaded at 0.05mm/sec and loads were measured using a 50lb load cell (Honeywell). Maximum load (N), ultimate load (N), yield load (N), stiffness (N/mm), energy (Nmm), yield displacement (mm), ultimate displacement (mm), and failure displacement (mm) were calculated from the load–displacement curves. Data from female-born mice were compared using two-way ANOVAs and corrected for multiple comparisons using Tukey’s multiple comparisons tests (Prism v9.0; Graphpad, LaJolla, CA).

## Results

Bone size was influenced by GAHT, with testosterone treatment with or without GnRHa administration leading to a significant decrease in femur length (Table 1: source of variation = T, p = 0.0011). Additionally, the shape of superstructures on long bones was influenced by GAHT (Figure 3A). Specifically, the width of the third trochanters was larger in mice treated with testosterone compared to control (without GnRHa or T administration) and GnRHa only treatment (Figure 3B, p = 0.0023). Mice treated with testosterone, with or without GnRHa, also had reduced cortical thickness adjacent to the third trochanter (Figure 3C).

**Table 1.** Biomechanical outcomes for femurs following 4-pt bending. * indicates that the effect of GnRHa treatment was significant, p= 0.0371; ** indicates that the effect of T treatment was significant; p = 0.0013; *** indicates group is significantly different than female mice with and without GnRHa or T, p <0.01; ^ indicates group is significantly different than no T or GnRHa females, p<0.01; ^ indicates group is significantly shorter compared to same GnRHa treatment with no T, p<0.01.

|  |  | <u>No GnRHa</u> |  |  | <u>GnRHa-treated</u> |  |  | <u>Male</u> |  |  |
| --- | --- | --- | --- | --- | --- | --- | --- | --- | --- | --- |
|  |  | Mean | 95% CI | N | Mean | 95% CI | N | Mean | 95% CI | N |
| Yield Load (N) | no T | 19.12 | 19.59 | 5 | 19.12 | 19.59 | 5 | 22.95 | 13.01 | 3 |
|  | with T | 18.78 | 18.57 | 6 | 16.98 | 12.51 | 5 |  |  |  |
| Ultimate load (N) | no T | 24.21 | 14.48 | 5 | 25.07 | 4.01 | 6 | 31.08 | 1.92 | 3 |
|  | with T | 25.99 | 9.18 | 6 | 25.62 | 4.83 | 5 |  |  |  |
| Stiffness (N/mm) | no T | 139.78 | 13.82 | 5 | 153.65 | 9.82 | 6 | 7.01 | 41.87 | 3 |
|  | with T | 151.03 | 10.40 | 6 | 148.35 | 7.33 | 5 |  |  |  |
| Energy (Nmm) | no T | 8.49 | 40.95 | 5 | 9.51 | 44.30 | 6 | 7.01 | 41.87 | 3 |
|  | with T | 9.27 | 26.15 | 6 | 9.54 | 22.94 | 5 |  |  |  |
| Yield displacement (mm)* | no T | 0.15 | 17.73 | 6 | 0.12 | 11.97 | 6 | 0.13 | 20.98 | 3 |
|  | with T | 0.14 | 24.03 | 6 | 0.13 | 7.83 | 5 |  |  |  |
| Ultimate displacement (mm) | no T | 0.27 | 5.04 | 5 | 0.25 | 6.89 | 6 | 0.25 | 5.93 | 3 |
|  | with T** | 0.29 | 7.25 | 6 | 0.29 | 7.60 | 5 |  |  |  |
| Femur length (mm) | no T | 15.08** | 2.36 | 6 | 15.34 | 1.55 | 6 | 15.76 | 2.08 | 3 |
|  | with T | 14.53****↓ | 1.93 | 6 | 14.76***↓ | 1.74 | 5 |  |  |  |

Treatment with testosterone, independent of GnRHa treatment, also led to significant gain in trabecular bone volume and number (Figure 4A and B; Table 2). This change was also associated with decreased trabecular spacing (Figure 4B). Testosterone treatment with or without GnRHa led to increased trabecular bone mineral density (BMD) with testosterone-only reaching comparable levels as trabecular BMD for age-matched male mice (Figure 4C). For most of the trabecular outcomes measured, female-born mice had significantly different trabecular bone properties compared to age-matched male mice, especially without T treatment (Table 2).However, with the inclusion of T treatment in female-born mice, trabecular properties were more similar to age-matched male mice independent of GnRHa treatment (Table 2). Unlike trabecular changes, we did not observe changes to cortical bone properties with GnRHa or T treatment in female mice (Table 3). However, most cortical bone properties, with the exception of cortical BMD, were significantly different between female-born mice with and without GnRHa or T treatment compared to age matched male mice (Table 3).

**Table 2.** Trabecular bone morphometry with and without GnRHA and T in female mice. If GnRHa was a source of variation (SoV) for morphometric outcomes, the associated p-values were noted in the morphometric column. If T was a SoV for morphometric outcomes, the signficance was noted in T-treatment columns as: * = p <0.05; *** = p <0.001; **** = p <0.0001. For outcomes in which male mice were significantly different than respective treatment groups, p-values are indicated in the treatment column as: * = p <0.05; *** = p <0.001; **** = p <0.0001.

|  |  | <u>No GnRHa</u> |  |  | <u>GnRHa-treated</u> |  |  | <u>Male</u> |  |  |
| --- | --- | --- | --- | --- | --- | --- | --- | --- | --- | --- |
|  |  | Mean | 95% CI | N | Mean | 95% CI | N | Mean | 95% CI | N |
| **Tb anisotropy MIL** | no T | 0.377 | 57.657 | 7 | 0.432 | 40.617 | 7 | 0.338 | 11.722 | 3 |
|  | with T | 0.362 | 32.750 | 7 | 0.375 | 25.403 | 7 |  |  |  |
| **Tb BV (mm<sup>3</sup>)**<br>GnRHa SoV,<br>p<0.001 | no T | 0.094** | 12.782 | 7 | 0.087*** | 25.118 | 7 | 0.289 | 2.277 | 3 |
|  | with T**** | 0.253 | 34.641 | 7 | 0.211 | 44.360 | 7 |  |  |  |
| **Tb TV (mm<sup>3</sup>)**<br>GnRHa SoV,<br>p<0.001 | no T | 2.535* | 9.778 | 7 | 2.632* | 3.943 | 7 | 3.398 | 5.992 | 3 |
|  | with T | 2.861 | 20.840 | 7 | 2.703 | 7.904 | 7 |  |  |  |
| **BV/TV**<br>GnRHa SoV<br>p=0.023 | no T | 0.037 | 11.319 | 7 | 0.033* | 22.453 | 7 | 0.085 | 6.736 | 3 |
|  | with T**** | 0.091 | 37.625 | 7 | 0.079 | 44.000 | 7 |  |  |  |
| **TbTh (mm)**<br>GnRHa SoV<br>p=0.0003 | no T | 0.041** | 9.088 | 7 | 0.043* | 7.902 | 7 | 0.049 | 2.462 | 3 |
|  | with T | 0.044* | 2.444 | 7 | 0.045 | 6.494 | 7 |  |  |  |
| **TbSp (mm)**<br>GnRHa SoV<br>p=0.298 | no T | 0.308 | 19.371 | 7 | 0.322* | 9.428 | 7 | 0.228 | 11.471 | 3 |
|  | with T**** | 0.201 | 17.221 | 7 | 0.225 | 17.940 | 7 |  |  |  |
| **TbN (1/mm)** | no T | 2.920 | 13.131 | 7 | 2.754* | 8.502 | 7 | 3.629 | 9.711 | 3 |
|  | with T**** | 4.139 | 12.645 | 7 | 3.766 | 14.186 | 7 |  |  |  |
| **Tb BMD (mg HA/cm<sup>3</sup>)**<br>GnRHa SoV<br>p=0.0007 | no T | 111.975** | 13.904 | 7 | 103.929** | 22.457 | 7 | 228.359 | 15.178 | 3 |
|  | with T**** | 224.018 | 27.406 | 7 | 185.092 | 30.431 | 7 |  |  |  |
| **Tb BMC (mg HA)** | no T | 0.285*** | 18.946 | 7 | 0.275**** | 25.015 | 7 | 0.772 | 11.490 | 3 |
|  | with T**** | 0.601 | 31.250 | 7 | 0.500 | 32.426 | 7 |  |  |  |
| GnRHa SoV<br>p<0.0001 |  |  |  |  |  |  |  |  |  |  |
| **Tb TMD**<br>(mg HA/cm<sup>3</sup>) | no T | 1146.584** | 12.827 | 7 | 1259.507 | 7.184 | 7 | 1522.848 | 3.775 | 3 |
|  | with T* | 1370.588** | 8.297 | 7 | 1281.477 | 8.818 | 7 |  |  |  |
| **Tb TMC**<br>(mg HA) | no T | 0.108*** | 21.266 | 7 | 0.110*** | 30.551 | 7 | 0.440 | 5.262 | 3 |
|  | with T**** | 0.355 | 41.321 | 7 | 0.277 | 51.834 | 7 |  |  |  |

**Table 3.**
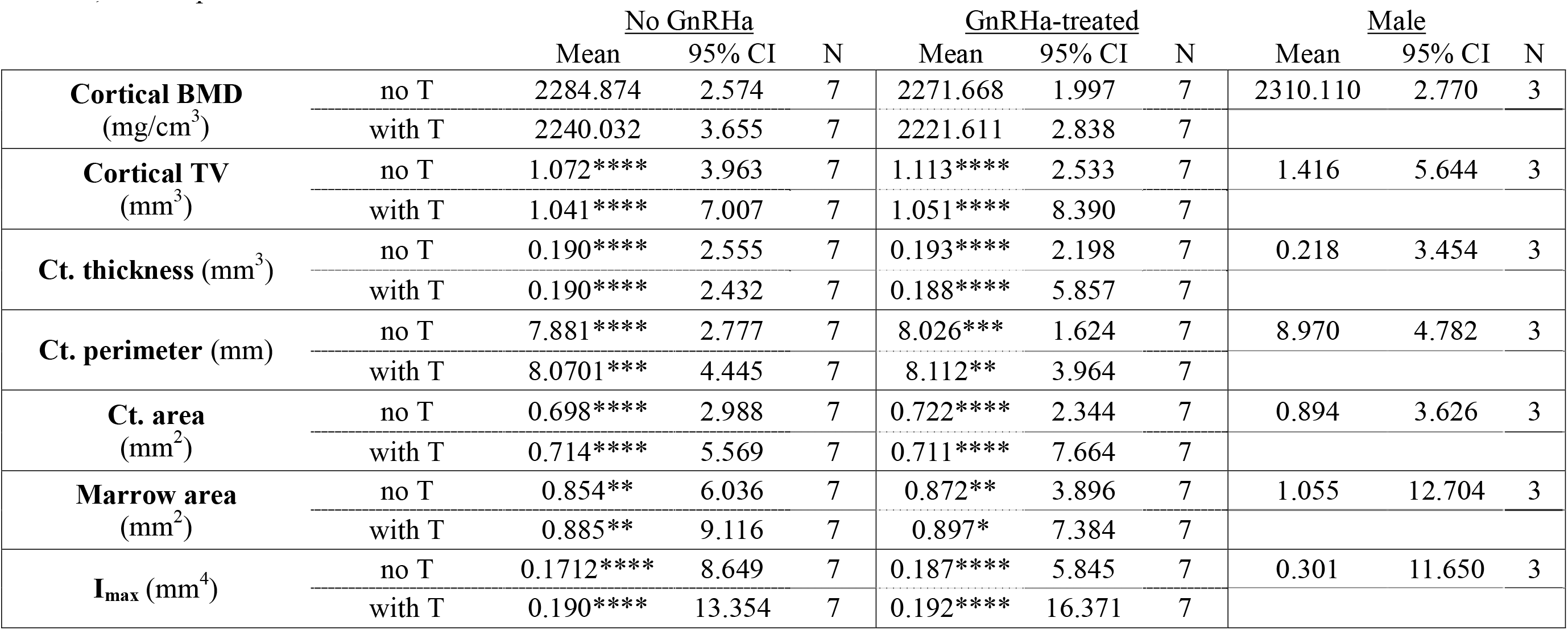
Cortical bone morphometry for female mice with and without GnRHa and T treatment. For outcomes in which male mice were significantly different than respective treatment groups, p-values are indicated in the treatment column as: * = p <0.05; ***=p<0.001;**** = p <0.0001.

**Figure 4.**
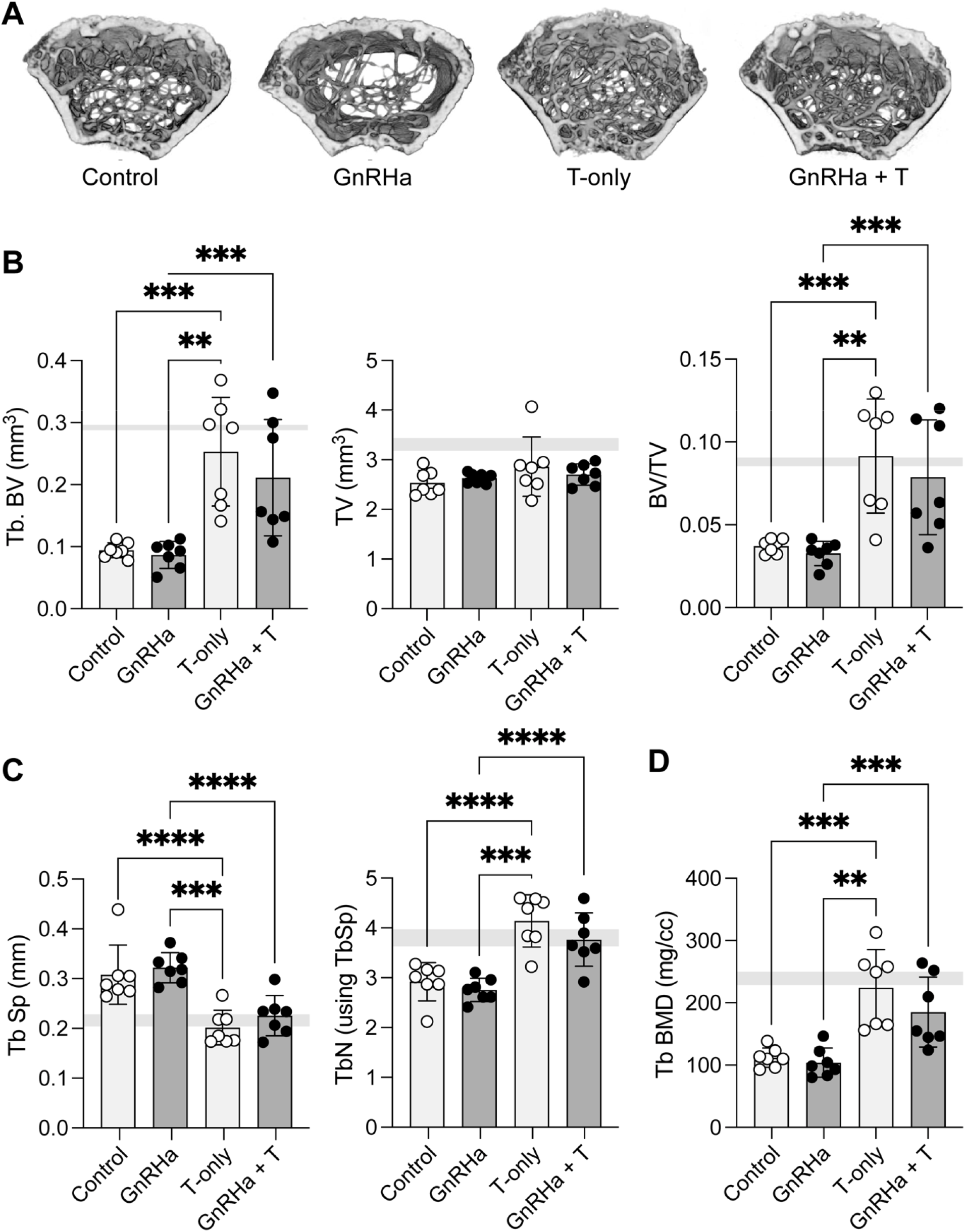
Trabecular bone parameters in mice prepubertally treated with gonadotropin-releasing hormone analogues (GnRHa) and supplemented with gender-affirming hormones (GAH). (A) 3D reconstruction of femoral trabecular bone from P89 female mice. A representative image from each group is depicted. (B) T-treated femurs showed a significant increase in trabecular bone volume and trabecular bone volume fraction (BV/TV). (C) Trabecular density and number were increased in femurs treated with T; (D) inversely T-treated groups displayed a lower trabecular separation compared to non-T treated mice. Trabecular Data are presented as biological replicates (individual dots) and mean +/-standard deviation. Gray horizontal bars represent range for age-matched male control mice, n=3. ** p < 0.01; ***p<0.001,**** p < 0.0001.

The effect of testosterone accounted for ∼40% of the variation in ultimate displacement (p=0.0013; Fig 5). T treatment accounted for 42.65% of the variation in femur length measured at time of biomechanical testing whereas GnRHa treatment accounted for 14.89% of the variation.

**Figure 5.**
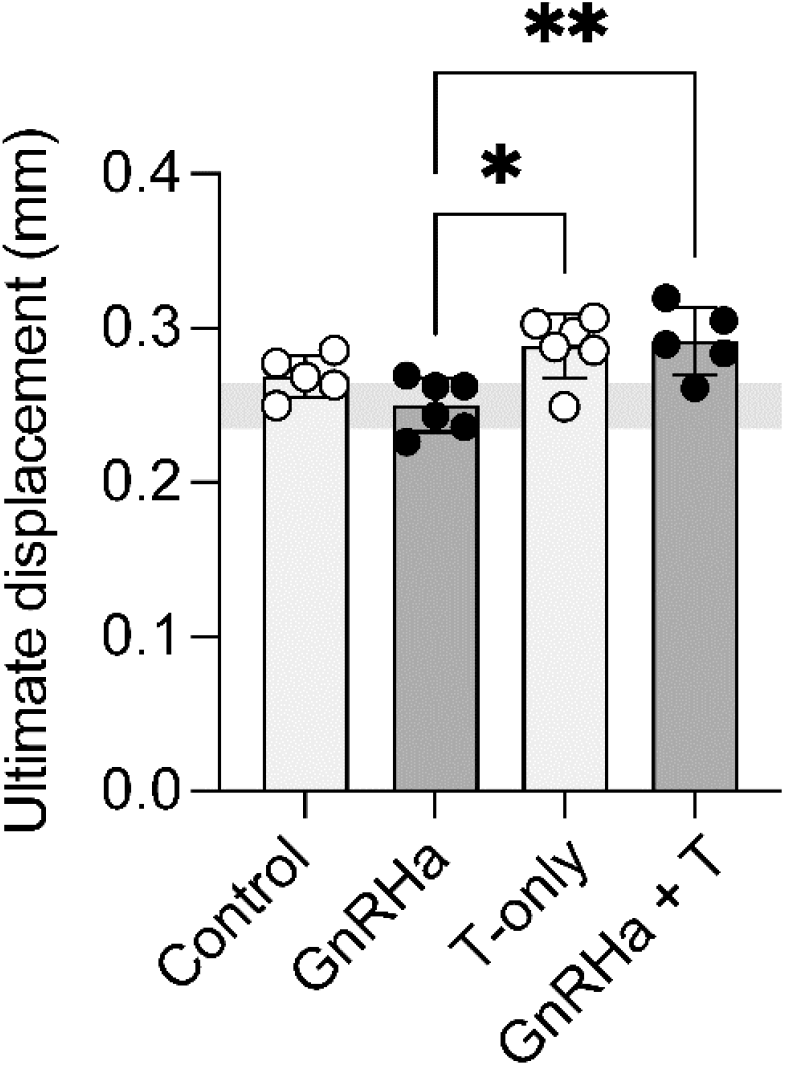
Ultimate displacement during fracture was significantly higher in female mice treated with T (with or without GnRHa) compared to GnRHa only mice. Data are presented as biological replicates (individual dots) and mean +/-standard deviation. Gray horizontal bars represent range for age-matched male control mice, n=3. * p = 0.0117; ** p < 0.01.

## Discussion

The effect of pubertal suppression in transgender and gender diverse youth followed immediately by sex steroid administration on bone growth and quality is a contemporary topic and important to investigate for the long-term health of these youth. Peak bone mass and body composition are strongly affected by sex steroids during puberty. High peak bone mass at skeletal maturity can predict a lower rate of fracture risk later in life (1–5). This is of special consideration for transgender and gender diverse youth who may elect to suppress puberty using GnRHa to delay or inhibit secondary sex characteristics. Estrogen plays a key role in bone regulation and growth, and conversion of testosterone to estrogen via aromatase or estrogen treatment in states of estrogen deficiencies can lead to increased bone remodeling. (22–26).

Estrogen treatment can accelerate skeletal maturity, as well as induce senescence of proliferating growth plate cells, leading to early fusion of the epiphysis (22). We believe that testosterone conversion via aromatase could be an explanation for our finding of a decrease in femur lengths in our T treated groups (Table 1). Peak longitudinal growth velocity is also correlated with levels of growth hormone (GH) and insulin-like growth factor 1 (IGF-1) (5). Estrogen has been shown to increase GH secretion in boys and girls during puberty, and this increase is observed earlier in girls (5). Coincidentally, testosterone has a similar effect on GH and IGF-1 but much of the effect of testosterone is mediated through its aromatization to estrogen (5,27–29). Future studies should explore how changes in GH and IGF-1 are associated with gender affirming care in the context of bone health.

GnRHa can also induce a hypoestrogenic state resulting in bone density loss and reduced bone health (6). Prior to the widespread use of estrogen “add-back” therapy, some studies showed immediate bone density loss ranging from 2%-6% after 6 months of GnRHa therapy, but this bone loss was recoverable (6). While the literature focused on the impact of gender-affirming therapy is growing, few have explored the effects of a hypoestrogenic period followed by subsequent GAH administration. Of note, in an adolescent mouse model of GAHT like ours, Dubois et al. showed that treatment of GnRHa led to fat mass accumulation, reduced lean mass and muscle strength, reduced of bone mass acquisition and bone strength, and increased bone marrow adiposity (8). Additionally, they found that T administration in prepubertal female mice can reverse the hypoestrogenic effects of GnRHa treatment (8). In the present study, we evaluated long bone (femur) quality in female-born mice subjected to peripubertal suppression followed by testosterone treatment. We found T-treatment with or without GnRHa during peripubertal growth had a strong effect on trabecular bone but only a marginal effect on cortical bone. Previous studies have demonstrated that bones with a well-developed trabecular network have increased bone strength (7–9). In our study, androgen administration led to a significant increase in trabecular bone density and number. We found that GnRHa treatment also resulted in longer femurs and, conversely, T administration led to shorter femurs, which could be explained by androgen-mediated acceleration of growth plate closure (30).

We also observed a visual difference in the shape of long bones, specifically focusing on the size of the third trochanter. With T treatment, we found the trochanter extended further from the centroid of the bone, and this may be caused by increased muscle contractility of the large gluteus maximus muscle. In young adults, testosterone-containing GAHT has been shown to improve knee extension and flexion strength (31). Further studies will begin to explore if there is an actual difference in trochanter size, including analysis of a transfeminine model (estrogen-containing GAHT).

In animals and in humans, bone mass measurements in adulthood can be used to assess both peak bone mass acquisition as well as monitor the subsequent decline caused by aging or gonadal insufficiency (23). Men with aromatase deficiency and women in menopause have lower bone mass, highlighting the impact of estrogen on bone maintenance (24). Conversely, women with polycystic ovarian syndrome (PCOS) and hyperandrogenism have increased trabecular BMD, and women with androgen insensitivity have lower BMD, suggesting that testosterone regulates BMD in women as well (24). In animal models, removal of gonadal sex hormone production in neonatal mice (e.g., ovariectomy or orchiectomy) can lead to restricted femoral bone growth during peripubertal development (24–26).

In summary this work demonstrates that delayed puberty in female-born mice does not significantly impair long bone quality or strength into young adulthood, and supplementation with T can lead to increased trabecular bone density. These gender-affirming therapies also influence the shape and length of long bones like the femur, which may correlate with changes in skeletal muscle function. These findings can help guide clinicians in their treatment of transgender and gender diverse adolescents by increasing the understanding of the effects of GAHT on long bones during a critical period of development.

## Acknowledgements

This work was supported by funding from the National Institutes of Health NIAMS (R01AR079367 to MLK and P30AR069620) and NICHD (R01 HD098233 to MM, AS and VP); the National Science Foundation (CAREER 1944448 to MLK). Schematics designed using Biorender.

